# Revisiting Demand Rules for Gene Regulation

**DOI:** 10.1101/014142

**Authors:** Mahendra Kumar Prajapat, Kirti Jain, Debika Choudhury, Gauri S. Choudhary, Supreet Saini

## Abstract

Starting with Savageau’s pioneering work from 1970s, here, we choose the simplest transcription network and ask: How does the cell choose a regulatory topology from the different available possibilities? We study the natural distribution of topologies at genome, systems, and micro-level in *E. coli* and perform stochastic simulations to help explain the differences in natural distributions. Analyzing regulation of amino acid biosynthesis and carbon utilization in *E. coli* and *B. subtilis*, we observe many deviations from the demand rules, and observe an alternate pattern emerging. Overall, our results indicate that choice of topology is drawn randomly from a pool of all networks which satisfy the kinetic requirements of the cell, as dictated by physiology. In short, simply, the cell picks “whatever works”.

## Introduction

A critical feature of all living organisms is the ability to tune behavior in response to stimuli [1-5]. The most widespread and well-understood mode of this tuning is transcription, which enables the cells to modulate gene expression in response to cues. Looking at the simplest transcription network, where a regulator R, controls expression of a target T - different possibilities emerge. Control of the target, might be via positive or negative regulation. When we consider the fact that most transcription factors in *E. coli* are also auto-regulators, six possible topologies emerge (Figure 1) [3, 4, 6-10]. In this study, we seek to answer the following question: Among all the available regulatory designs, how does a cell pick one to control target expression?

**Figure 1.**
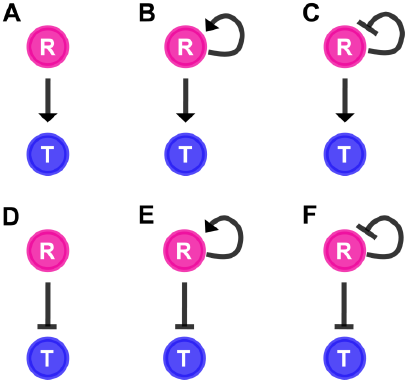
Topologies between a regulator R (pink) and a target, T (blue) In topologies A, B, and C, in presence of the appropriate environmental or cellular signal, target T is controlled positively by the regulator. In contrast, target in topologies D, E, and F is repressed by regulator, and the repression is relieved under appropriate conditions.

In a series of papers in the 1970s, Savageau proposed “demand rules for gene regulation” [11-16], according to which, a target T is positively regulated (Figure 1A-C) if, in the organism’s natural habitat, T is required for a high fraction of time. On the other hand, if the target is only required sporadically, it tends to be regulated negatively (Figure 1D-F) [12, 13]. Evidence was provided in the shape of conformity in regulation of sugar utilization enzymes in *E. coli* with the demand rules [11, 12]. In 2006, Alon et. al. provided a functional explanation for demand rules [17]. They argued that positive regulation for a frequently needed target T ensured erroneous binding of other transcription factors to the promoter was minimized. Alon et. al. demonstrate and propose, in a later report [18], that such an approach for gene regulation acts as an insulator of the promoter regions, preventing erroneous transcription.

However, demand rules raise a few interesting questions. Active control, as proposed by the demand rules, will increase the demand of regulators in the cell. The cost associated with production of additional regulators for control is likely detrimental for cellular growth [19-21]. In addition, demand rules seem contrary to the concept of genetic robustness, which focuses on loss of fitness due to mutations acquired by an individual [22]. How then do we reconcile these seemingly opposite logics? In a 2009 report, Hwa et. al. have, via a theoretical framework, demonstrated that the choice of mode of gene regulation could be biased for or against demand rules, and is dictated by population size and the time scale of environmental evolution [23]. Their framework remains to be experimentally tested though. An alternate approach can be to examine response of different topologies to cues. The response can be quantified in terms of parameters like time of response, response to noise, and cost of control [3, 24-28]. However, questions like whether, over physiologically relevant range of biochemical parameter values, there are inherent qualitative differences in the response that can be generated by different topologies remain unanswered.

In this work, we perform simulations of the simplest transcriptional network (Figure 1), and compare our results with the natural distribution of regulatory interactions among topologies in *E. coli*. We revisit some of the results proposed by Savageau and study in detail the control of sugar utilization and amino acid biosynthesis in *E. coli*. Finally, we characterize the role of control cost in dictating fitness of a cell. Put together, our results indicate that, contrary to demand rules, choice of a particular topology for gene expression control is likely chosen randomly from all available topologies which satisfy the dynamic demands of physiology associated with a cellular function.

## Results

### At the global scale, E. coli chooses topologies differentially for control of gene expression

To understand the “logic” behind choice of topology for gene expression control, we enumerated all regulatory interactions in *E. coli*, and classified them in one of the six topologies in (Figure 1) [29]. We note that there is no qualitative difference in the number of interactions which are controlled via positive (∼49.6%) or negative regulation (∼50.4%) (Figure 2A). Including global regulators (and their regulons) in the enumeration yields similar results (Figure S1(A)). We also repeated the analysis by defining an interaction as a regulator R controlling a promoter (instead of all genes in an operon) (Figure S1(B-C)), and the analysis exhibits that there is no significant bias in *E. coli* choosing positive or negative regulation preferentially.

**Figure 2.**
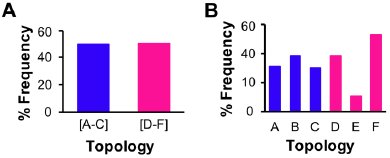
Frequency distribution of R - T topologies in *E. coli*. **(A)** All regulator-target interactions frequency in *E. coli* among activator- (topologies A, B, C) and repressor-based (topologies D, E, F) interactions. **(B)** Distributions of all R-T interactions among the six distinct topologies in *E. coli*.

However, the R-T frequency distribution changes qualitatively when we analyze the number of interactions in each of the six topologies in (Figure 1). As represented in (Figure 2B), among the six topologies, F is over-represented. This is followed by topologies A, B, C, and D, with no statistically significant difference between them. Last, topology E is the least represented (∼5% of all interactions). We observe the same general trend when we define one interaction as regulator R controlling a promoter, instead of a gene (Figure S2A). On including the global regulators from the analysis, a slightly different picture emerges, where topologies C and F are the most represented (as most global regulators auto-regulate themselves), followed by topologies B, D, A, and E (which is again under represented) (Figure S2(B-C)).

Overall, our analysis suggests that *E. coli* prefers certain regulatory arrangements over others. What are the factors that dictate this choice? Various possibilities exist, including, demand rules [11-16], error-minimization [17], or minimizing cost of control [19-21]. To understand the differences between the distribution of the six topologies, we study and analyze their distribution at two different scales. At the first level, we analyze frequency distribution of regulatory arrangements at a systems level, where a system is defined as sum of all interactions which serve the cell towards a common function. At the second level, we analyze in detail the demand and corresponding regulatory design at a micro level, specifically those related to amino acid biosynthesis and carbon source utilization.

### Differential choice of topology at a systems scale

To analyze frequency distribution of topologies in further detail, we separated R and T interactions in *E. coli* into six functional sub-groups: (a) amino acid transport and metabolism, (b) sugar metabolism and energy production, (c) coenzyme transport and metabolism, (d) inorganic transport and metabolism, (e) cell division and nucleotide metabolism, and (f) stress response (Figure 3). In five of the six classifications, distribution of number of interactions among A-C and D-F is statistically identical to the distribution observed at the genome scale in *E. coli*. Moreover, when analyzed individually, we note that in each of the six classifications, topology E is under-represented, and topology F is over-represented. In addition, the frequency distributions in all six groups is statistically similar to the one observed in nature (Figure 2B). We revisit the reason and nature of this distribution later in this manuscript. It is not surprising that topology F is over-represented in nature. Negative auto-regulation is known to speed up response in cellular systems [9, 25], and both topologies C and F possess that architecture. However, subtle differences exist. While topology C speeds up response when the system transitions from an OFF to an ON state, topology F speeds up cellular response in transition from ON to OFF state. Could other similar dynamic criteria explain the differential use of topologies in *E. coli*? To help answer this question, stochastic simulations of the simplest regulatory network between a regulator R and target T were performed (Figure 1).

**Figure 3.**
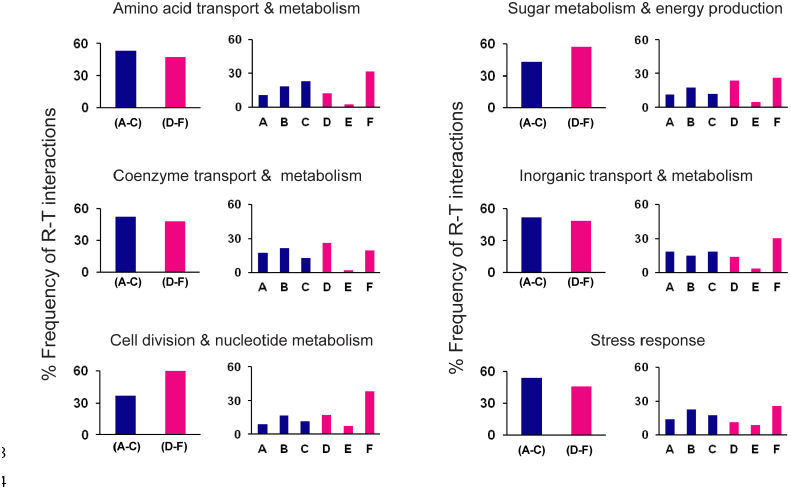
Division of R-T interactions among six functional groups. For each group, frequency of activator-based topologies roughly equals repressor-based topologies. Among individual topologies, F is over-represented and E under-represented in all six functional groups.

### Simulations to quantify performance of networks across topologies

We define a list of factors that best define performance of a regulatory circuit. These include (a) steady state target expression, (b) time of response, (c) control cost, (d) cell-to-cell variation, (e) spread of gene expression, and (f) ability to be effectively switched ON and OFF (see methods for more details). We hypothesize that these six indicators define performance of a genetic network. Network dynamics are dictated by the values of the associated biochemical parameters. To account for biases introduced by parameters, we simulated about 100,000 networks (∼16,000 networks in each topology). The choice of parameters for these networks was taken from a uniform spread from a range. Each network was simulated for 500 cells, and transition from OFF to ON and ON to OFF tracked.

Because of our choice of parameters from a parameter space, many networks are “dead” (steady state target expression less than one); have infinite cost; or have physiologically unviable dynamics. For analysis, we considered only those networks which express target T and are able to effectively switch ON/OFF. In addition, we impose limits on the time of activation (& deactivation) and control cost. Placing these constraints allows us to define a “Performance Box”. In rest of the article, we only consider networks which lie within this “Performance Box”. A frequency count of the networks with positive and negative control of T shows that the two are identical (Figure 4A), consistent with the natural distribution in *E. coli*.

**Figure 4.**
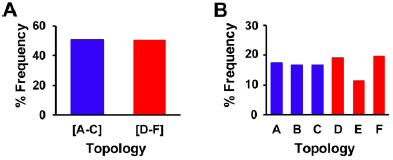
Frequency distribution of R - T topologies from simulations. **(A)** Percent networks belonging to the activator-and repressor-based topologies in the “Performance Box” and **(B)** Percent networks belonging to each of six topologies in the “Performance Box”.

Frequency distribution of the six topologies shows that, just as in *E. coli*, topology E is underrepresented, and topology F, most represented in the “Performance Box” (Figure 4B). Hence, our simulations suggest, and we speculate that distribution of a topology in a cell is proportional to the frequency of the topology in the “Performance Box”. We note, however, that there are differences in the distribution of the networks among the six topologies between our computational results and the *E. coli* distribution. We hypothesize that these differences in distributions are due to the inherent differences in the dynamic features of the six topologies.

#### Dynamics of activation and deactivation vary across the six topologies

Analysis of time of activation and deactivation indicate that subtle differences exist in the T-t50 space covered by topologies A-C and D-F (Figure 5A-B). As shown in (Figure 5A), topologies D-F are more suited for slow activation of targets with small steady state levels. In the OFF state (Figure 5B), we note that physiological functions with preference for smaller steady state T values and higher deactivation times would have a greater chance to be represented by topologies A-C. Qualitative differences exist in performance associated with each of the six topologies (Figure S3). In the ON state, topology C and F exhibit the widest range of steady state T and activation time. On the other hand, topology E is the most constrained in terms of the possible values of T. We speculate that this is one of the reasons of over-representation of topology F, and under-representation of topology E in natural circuits.

**Figure 5.**
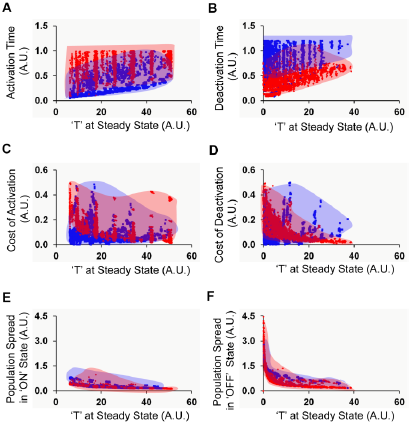
Dynamic performance of activator-and repressor-based topologies. **(A-B)** Steady state T levels and time of activation/deactivation for all six topologies. A dot on the plot represents each network. **(C-D)** Steady state T and cost of control phase plane for the six topologies. **(E-F)** Steady state T and spread among the 500 cells at steady state in the topologies. Activator-based topologies are represented in blue, and repressor-based topologies in red. Shaded regions indicate the region covered by the activator-and repressor-based topologies. The left panel represents networks in ON state. Right panel represents OFF state.

#### Cost of control of expression

(Figure 5C-D) show that while there are large T-cost space regions which both groups of topologies can exhibit, only topologies A-C can control expression when T is small. Moreover, this control is exhibited at relatively small control cost. Similarly, when switched OFF, topologies A-C are better able to switch expression off when the desired T values are small. In addition, on an average, topologies A-C are able to provide control with a smaller drop in T when cells are moved from ON to OFF, the physiological significance of which remains unknown. Comparison of each topology is presented in (Figure S4). During activation, the control cost in topologies A-E decreases with increase in T. However, in topology F, the decrease is super-linear, and the resulting fall in cost much more rapid. This difference likely places topology F at an advantage when high expression of target T is required. Qualitatively, from the pattern of (Figure S4), we note that topologies A-C behave identically, whereas D and E behave differently. During deactivation, all topologies (except C) offer a similar pattern of deactivation dynamics and steady state T. We also note, that for topology F, T (in OFF state) is a considerable fraction of T in ON state. This suggests that topology F is best suited for genes which require small changes in expression levels in different conditions.

#### Cell to cell variation

The stochastic nature of gene expression leads to heterogeneity at a single-cell resolution. Our results show that across the two groups of topologies, there is little qualitative difference in behavior (Figure 5E-F). Among individual topologies, two key differences exist. First, in the ON state, topology F is able to exhibit behavior with the widest range of spread in expression of T. Second, in OFF state, topologies with negative control, all exhibit greater spread in the steady state T values as compared to topologies A-C. Cell to cell variation is likely key for cells to survive in uncertain conditions. This is likely another reason for presence of higher than random frequency of F networks in nature (Figure S5).

#### Switchability

For optimal physiology, *E. coli* would need to control expression of a large number of genes. However, the control and tuning of each gene in the OFF or ON state will be unique. Hence, each physiological role would require a topology most suited to provide appropriate control. From our analysis (Figure 6), we note that in this respect, topology F offers the widest ratio of steady state T in ON and OFF conditions, across a large range of T values. Networks in topologies A-C offer a qualitatively similar and a very limited response dynamics in this regard. In addition, topology E offers the most limited response in terms of steady state T values, and hence is likely least suited for most cellular functions.

**Figure 6.**
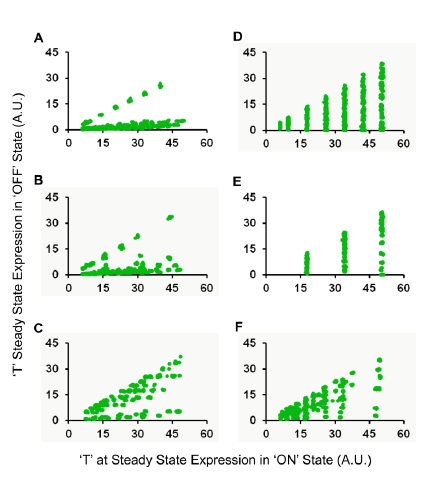
“Switchabililty” of networks. Steady state T in the ON (a-axis) vs. the OFF condition (y-axis) for all six topologies.

### Revisiting Demand Rules for Gene Regulation

To test and apply the demand rules in determining the choice of regulatory topology, we perform two analyses at a micro scale. In the first, we study amino acid biosynthesis in *E. coli*, primarily present in mammalian intestine, and with ability to synthesize all 20 amino acids. However, because of unequal presence in the intestine, not all amino acids are required equally by the bacterium [30, 31]. The demand for amino acids is further biased by the number of codons for each amino acid in the *E. coli* genome (**Sheet S1, Excel**). The actual availability of amino acids was therefore calculated as availability normalized with demand for an amino acid in the *E. coli* genome. In our analysis, we identified the biosynthesis pathway(s) which are uniquely dedicated to synthesis of a particular amino acid only [32], analyzed regulation of each gene in the pathway(s), and classified regulation as positively or negatively regulated topologies.

Amino acid biosynthesis and transport are cellular functions with inversely related demand. For instance, if an amino acid is not present in the surroundings (resulting in low demand for transporters), the biosynthetic demand would be high, and vice versa. In (Figure 7A), the x-axis represents amino acids in increasing availability in *E. coli* habitat, while the y-axis gives the fraction of all regulatory interactions, controlling biosynthesis and transport of that amino acid, belonging to topologies A-C. Our analysis shows that regulation of both biosynthesis and transport exhibit a statistically insignificant correlation with increasing availability. This is contrary to the demand rules. Adherence to the demand rules would have meant that transporters of abundant amino acids are regulated by A-C topologies, and biosynthetic genes for such amino acids are primarily regulated by D-F topologies. The reverse would have held true for scarcely available amino acids. The same results hold on including interactions involving global regulators (Figure S6A). (Figure S6B-C) show the data when amino acid demand is not normalized with the number of codons in the genome – both the results show statistically insignificant relationship against demand rules. Additionally, we performed a similar analysis for the soil bacterium *B. subtilis*, and found no correlation between choice of topology and availability of an amino acid in the surroundings [33, 34] (Figure S7).

**Figure 7.**
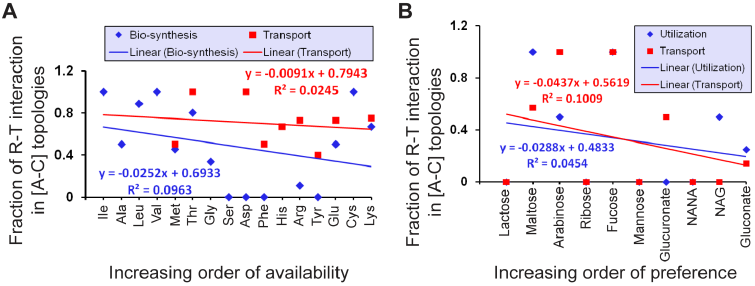
Correlation between demand and activator-based control. **(A)** X-axis represents amino acids in normalized increasing availability, and y-axis represents fraction of regulatory interaction in the activator-based topologies for each amino acid biosynthesis regulon (blue) and its transport (red). **(B)** X-axis represents carbon sources in increasing order of preference to *E. coli* in the intestine, and y-axis, the fraction of regulatory interactions controlling expression of metabolic genes involved in utilization (blue) of each sugar and its transport (red) and belonging to activator-based topologies.

The demand for proteins encoded by amino acid biosynthesis genes is inversely linked with the presence of amino acid in the surrounding environment. Hence, we would expect that expression pattern (and demand) of biosynthetic genes and amino acid transporters is linked inversely. However, performing a similar analysis on amino acid transporter gene regulation in *E. coli* demonstrates a lack of correlation between demand for the gene product and choice of topology (Figure 7A).

In the second example, we focus on metabolism of sugars preferred by *E. coli* in its natural habitat *[35]*. Based on their abundance, we obtained the relative demand for carbohydrates in the intestine. For our analysis, we only considered part of metabolism which deals exclusively with a particular carbon source. The genes encoding the respective enzymes and their regulators were studied, and classified into activator-or repressor-based topologies. Our results indicate that regulation of enzymes involved in carbon utilization is independent of the availability of the sugar (Figure 7B). A similar statistically insignificant result was obtained on including global regulators (Figure S8). In case of carbon utilization, the expression of transporter genes is positively correlated with expression of genes involved in catabolism. However, our analysis shows that the choice of topology does not seem linked with availability of the carbon source (Figures 7B and S8). Similar analysis was performed for carbon utilization in *B. subtilis* and no statistical correlation was observed between choice of topology and demand for product of gene of interest [33, 34](Figure S9).

Overall, our results indicate deviations from Savageau’s demand rules. A major difference in our and Savageau’s analysis is that we consider all regulatory interactions controlling cellular functions, while Savageau’s work only accounted for the key regulator involved in a particular cellular process, for example, AraC for arabinose catabolism [11, 13]. Another key difference lies in the fact that our analysis takes into account auto-regulation of regulatory proteins. This is likely extremely important in dictating the choice of topology, as seen by differences in topologies D, E, and F in *E. coli*.

### Cost of control places a growth burden on the cell

Cellular growth is hindered by production of unnecessary proteins [19-21, 36], and as a result, regulation has a fitness effect [37, 38]. To test this, we performed competition assays between genetically identical strains with the only difference that one strain was producing GFP. Our analysis with the *rob* promoter in *E. coli* demonstrates that the cost of additional GFP places a growth burden on the cell (Figure 8). Similar trends for P*araBAD* and P*marRAB* promoters were also observed. This suggests that the advantage of preventing erroneous transcription by adhering to the demand rules must offset the disadvantage of additional control cost, for demand rules to prevail.

As a speculative test of the demand rules, we performed long-term experiments where we fed a arabinose to *E. coli* for 3000 generations. In parallel, we fed glucose with limited arabinose to the culture. The culture grown on arabinose had high demand for arabinose utilization genes, whereas the culture with small amounts of arabinose would only express from *araBAD* operon when out of glucose - thus creating differential demand for the *araBAD* gene products. AraC is known to be a dual regulator of the *araBAD* operon, acting as a repressor in absence of arabinose, and activating expression when bound with arabinose. This dual regulation can be observed experimentally in wild type *E. coli*. On altering demand for the *araBAD* gene products, the dual regulation can still be observed in both (with high, and low demand for *araBAD* gene products) the strains (Figure 9A-B). In a relatively short span of 3000 generations, no switch in mode of regulation was observed, though the absolute levels of expression were different in the two strains, and had evolved from the parent wild-type *E. coli*.

**Figure 8.**
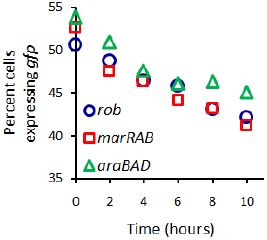
Competition between *E. coli* strains exhibits a fitness effect of production of an additional protein as compared to the control.

**Figure 9.**
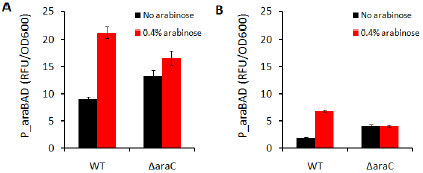
Long term experiment to track changes, if any, in mode of regulation. *P*_*araBAD*_ expression in **(A)** strain grown in 0.4% arabinose for 3000 generations. **(B)** strain grown in 0.35% glucose and 0.05% arabinose for 3000 generations. WT strain refers to the evolved strain after 3000 generations in a particular condition. Δ*araC* strain refers to the mutant created by knocking out *araC* from the parent evolved strain.

## Discussion

Transcription networks are of interest from a number of perspectives like structure, topology, dynamics, and evolution [3, 24, 39-42]. Despite significant effort in trying to understand dynamic features of topologies – an open question remains. Why does a cell choose a particular topology over the others?

Demand rules provide an insight into this question. However, our analysis reveals that the agreement with demand rules is rather limited. What then could be the additional determinants? Dynamically, as our analysis shows, there are several subtle differences across topologies, and it could be these differences which dictate choice. Our simulations and comparisons with *E. coli* distribution suggests that a network is perhaps randomly picked out from a group that satisfies the demands of physiology. Or simply, the cell picks “whatever works”.

In a 2009 study, Hwa et. al. demonstrated that different modes of regulation lead to qualitatively different patterns of protein levels when cells are grown in conditions supporting different growth rates [43]. They demonstrate that constitutively and positively controlled genes exhibit a decrease in steady state with growth rate, negatively regulated genes can exhibit a weakly negative or a strongly positive correlation between protein levels and growth rate. Could additional considerations like maintenance of protein levels at a constant levels, independent of growth rate, be a selective force for certain physiological roles?

In terms of what we can explore, our simulations very rapidly approach saturation in as we begin to increase the complexity of networks. In addition, combinatorial inputs of multiple regulators into one promoter remain unanswered and unexplored. These additional interactions would make the possible range of dynamic behaviour much more complex and richer, but at the same time computationally intractable.

## Experimental Procedure and Mathematical Analysis

### Regulatory interactions in E. coli

The R-T interaction dataset for the transcription regulatory networks was acquired from RegulonDB [29]. There are 197 transcription factors reported in RegulonDB, of which, seven are listed twice, individually, as well as in dimeric form with another protein. We have considered 190 unique regulators for our study and their interactions with targets have been classified in two ways; (i) between regulator protein and target gene and (ii) between regulator protein and promoter (all genes in an operon). This resulted in 4970 interactions in Regulator-Target gene classification (**Sheet S2 in Excel**) and 2139 interactions in Regulator-Promoter classification (**Sheet S3 in Excel**). Out of 190 transcription factors, seven are global regulators (CRP, H-NS, Lrp, IHF, ArcA, Fis, and FNR) and control around 51% of all genes in *E. coli* [44]. Excluding interactions of global regulators, there are 2625 interactions in regulator-target gene class and 1176 interactions in regulator-promoter class. Multiple transcription factors feeding into a promoter were categorized into more than one topology, depending on the nature of interaction of the target gene with each interacting transcription factor.

### Distribution of R-T interactions among six topologies

On the basis of the specific roles in cellular physiology, the interactions were further classified into six functional sub-groups, as reported in EcoCyc [32]. For each functional sub-group, all the involved target genes were identified and distributed among six topologies. Biosynthesis pathways for amino acids and degradation pathways for carbohydrate were studied further in detail.

#### Biosynthesis pathways of amino acids

For biosynthesis of an amino acid, we considered regulation of only target genes which play a role in biosynthesis (**Sheet 4 in Excel**) of that particular amino acid and its transport (**Sheet 5 in Excel**) only.

#### Frequency of occurrence of amino acids

Occurrence of each of the 20 amino acids from coding region of *E. coli* DH10β genome was calculated to calculate the relative demand of all amino acids in *E. coli* (**Sheet 1 in Excel**).

#### Sugar utilization

Genes encoding for enzymes involved in metabolism of a particular carbon source until the metabolic branch merges with another in the network were considered in our analysis (**Sheet 6 in Excel**). The interactions between the identified genes and their regulators (R-T) have been distributed across the defined topologies. In addition, genes involved in transport of sugars were also analysed in same way (**Sheet 7 in Excel**).

### Mathematical analysis of the six topologies

Mathematical model for each topology was formulated by writing ordinary differential equations, and simulating stochastically. The two differential equations for each topology describe the rate of change in regulator, R and target, T, as described in Supplement text.

#### Definition of parameter space

To analyze the six topologies, networks were generated with different biochemical parameters. To choose parameter values and range, physiologically observed values of all parameters was analyzed and the resultant space called parameter space (Figure S10A). From parameter space, the red region represents commonly observed values reported in literature [41, 45-50], biased towards exhibiting limited diversity in dynamics. In this work, we chose the “unbiased region” (blue) from parameter space to explore all possible dynamics [41, 45-50].

#### Generation of networks from a topology

The model of topologies B, C, E, and F consists of seven parameters whereas of each from A and D consists of five parameters. We generated 16807 (7^5^) networks for topology A & D and 16384 (4^7^) networks for topologies B, C, E, and F. Such an approach was recently adopted by Ma and co-workers in the context of analysis of adaptation in biochemical networks [51]. We also performed simulations with several other parameter distributions across the range of values. However, different parameter ranges do not qualitatively affect our analysis.

#### Calculation of performance indicators

Each network was simulated using Gillespie algorithm to account for stochasticity [52, 53]. Dynamic simulation of each network in both transitions from OFF to ON and from ON to OFF state was performed in 500 cells. The detailed flowchart of simulation is described in (**Figure S10B**). The dynamics of each network was recorded and performance indicators for each network were calculated as described in the Supplement.

#### Definition of “Performance Box”

A box in the performance indicator space defining networks with robust response, low cost, and fast dynamics was named as “performance box”. Networks with steady state expression of T ≥ 6 (A.U.) in ON state, with a minimum switchability factor of 1.3; activation time (t_50_) ≤ 1 (A.U.) and deactivation time ≤ 1.2 (A.U.); and cost of activation and cost of deactivation ≤ 0.5 (A.U.) were considered to define the outer edges of the “Performance Box”. Networks outside of the performance box were assumed to be more costly, exhibiting minimal expression of target protein T, or/and slow responding networks, and hence excluded from our analysis. The precise definition of the edges of the performance box presented here represents a general trend.

### Cost experiment using Flow Cytometry

*E. coli* containing a *rob* promoter fusion with *gfp* integrated at the λ site (plasmid pLA2) on the chromosome [54, 55] was grown overnight in LB media with kanamycin 25μg/ml in at 37°C with shaking. *E. coli* with pLA2-gfp (without the *rob* promtoer) was used as a control. Both cultures were grown overnight, and then sub-cultured in 1:500 dilution each in LB media with kanamycin 25μg/ml in the same tube and grown at 37°C with shaking. Samples were collected at various times and then stored in PBS (containing 34μg/ml chloramphenicol). All the samples were kept on ice in dark condition. The samples were then analyzed with BD FACS Aria SORP to get relative frequency. The choice of *rob* promoter was based on its constitutive expression [56]. Competition experiments were done with the *ParaBAD* promoter and the *PmarRAB* promoter, and similar results observed when the inducers for the two systems (arabinose and salicylic acid, respectively) were added to the media [57, 58].

### Evolutionary experiments

*E. coli* grown overnight in LB at 37°C with shaking was sub-cultured (1:100) in tubes containing 1ml M9 media, 1% casamino acids and a sugar source. The tubes either contained 0.4% arabinose or 0.35% glucose with 0.05% arabinose. The cultures were grown for 24 hours at 37°C, and propagated daily by sub-culturing 1:100 into fresh M9 media containing respective sugars. The last strains from each lineage was transformed with plasmid based promoter fusions of arabinose metabolic genes (*araB* (PEC3876-98156236)), from Thermo Scientific *E. coli* promoter collection (PEC3877). Fluorescence (488/525nm) and absorbance (600nm) values were measured in a Tecan microplate reader (Infinite M200 PRO).

## Acknowledgements

The work was funded by the Innovative Young Biotechnologist Award (IYBA), Department of Biotechnology, Government of India.

